# Cell type deconvolution of methylated cell-free DNA at the resolution of individual reads

**DOI:** 10.1101/2022.09.30.510300

**Authors:** Pia Keukeleire, Stavros Makrodimitris, Marcel Reinders

**Affiliations:** Delft Bioinformatics Lab, Delft University of Technology, Delft, the Netherlands; Translational Cancer Genomics group, Erasmus Medical Center, Rotterdam, the Netherlands; Leiden Computational Biology Center, Leiden University Medical Center, Leiden, the Netherlands

## Abstract

Cell-free DNA (cfDNA) are DNA fragments originating from dying cells that are detectable in bodily fluids, such as the plasma. Accelerated cell death, for example caused by disease, induces an elevated concentration of cfDNA. As a result, determining the cell type origins of cfDNA molecules can provide information about an individual’s health. In this work, we aim to increase the sensitivity of methylation-based cell type deconvolution by adapting an existing method, CelFiE, which uses the methylation beta values of individual CpG sites to estimate cell type proportions. Our new method, CelFEER, instead differentiates cell types by the average methylation values within individual reads. We additionally improved the originally reported performance of CelFiE by using a new approach for finding marker regions that are differentially methylated between cell types. This approach compares the methylation values over 500 bp regions instead of at single CpG sites and solely takes hypomethylated regions into account. We show that CelFEER estimates cell type proportions with a higher correlation (*r*^2^ = 0.94*±*0.04) than CelFiE (*r*^2^ = 0.86*±* 0.09) on simulated mixtures of cell types. Moreover, we found that it can find a significant difference between the skeletal muscle cfDNA fraction in four ALS patients and four healthy controls.

## INTRODUCTION

As cells die, short DNA fragments are released into the bloodstream, which are collectively called cell-free DNA (cfDNA). The cfDNA in plasma is mostly composed of DNA molecules originating from blood cells (1). However, cells in diseased tissues die more rapidly, causing diseased tissues to release cfDNA at a faster rate. The discovery of traces of such disease-derived cell types in cfDNA provides a minimally invasive alternative for tissue biopsies, and is thus frequently referred to as a liquid biopsy (2). Commonly researched applications of liquid biopsies are prenatal testing, organ transplant monitoring and tumor discovery and monitoring (3). In all of these applications, however, we know the cell type of interest in advance. Cell type deconvolution, on the other hand, aims to give the full composition of the cell types of circulating cfDNA. Example use cases in which this type of analysis is especially desirable is finding tumor locations in patients with a cancer of unknown primary (4) and detecting treatment side-effects. Additionally, characterizing changes in cell type proportions is helpful in understanding disease development and progression (5).

One method for characterizing the cell type origins of cfDNA is the analysis of methylation signatures. Methylation occurs when a methyl-group is added to the fifth carbon of cytosines (5mC), often with the purpose of silencing gene transcription (6). This process happens mostly in the context of CpG sites, and usually over regions spanning multiple CpG sites (2). Adjacent CpG sites have been found to correlate highly in methylation status (7). Because the silencing of gene transcription often happens in a cell type-specific manner, these methylation signatures have been found to reveal the cell type origins of cfDNA (3).

Traditionally, cell type deconvolution methods calculate the average methylation of all sequencing reads per CpG site, and use these averages as model input (8, 9, 10). These averages are often referred to as *β* values. Although these methods usually do take the correlation between nearby CpG sites into account by averaging over the *β* values in a region, the value at each individual CpG site is assumed to be independent.

In a similar problem setting, namely tumor fraction estimation, Li et al. devised an approach to better incorporate the correlation between sites (11). Their method, named CancerDetector, calculates the average methylation per individual sequencing read instead of the average methylation per CpG site. They showed that this method outperforms a similar previous method that uses *β* values (9), having higher sensitivity and specificity (11). Figure 1b illustrates how rare cell types can be more sensitively detected using read averages than using *β* values (11). In this figure, the tumor-derived read makes up 25% of the cfDNA, whereas such rare cell types are far less prevalent in biological data. Since it is essential that our method can deconvolve lowly abundant cell types, *β* values might not be appropriate.

**Figure 1.**
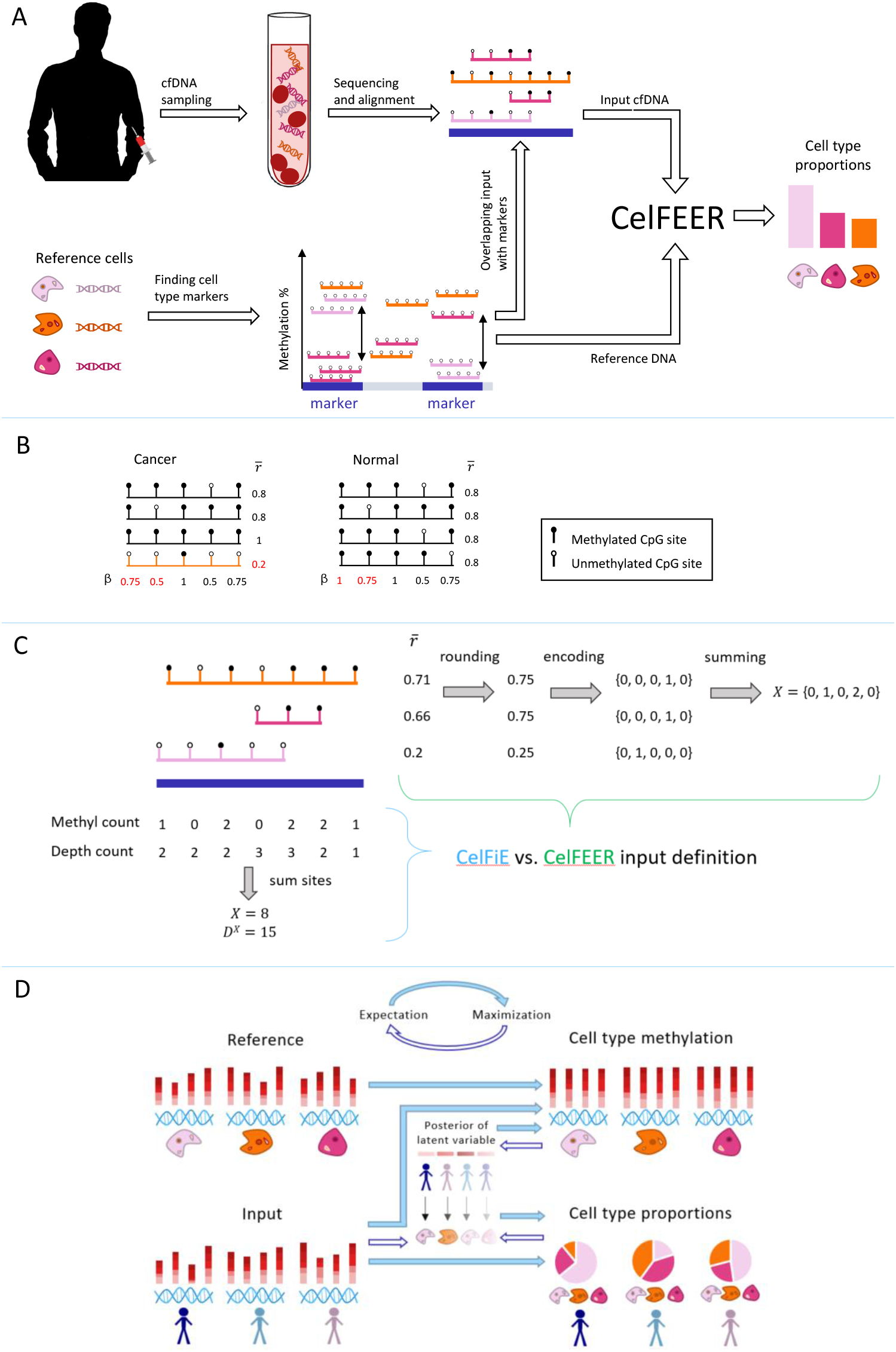
**(A)** Workflow of cell type deconvolution with CelFEER. Sequenced and aligned cfDNA fragments are intersected with cell type marker regions in the genome that are found using reference cell type data. The reference cell data and the cfDNA input data are used as model input for CelFEER, which subsequently outputs the estimated cell type proportions in the cfDNA. **(B)** Toy example illustrating how a tumor-derived read (in orange) can be distinguished from other reads more easily by comparing read averages 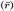 instead of CpG site averages (*β*). Values in red are differential between the cancer and normal sample. **(C)** Formatting of the input for CelFiE (bottom left) and CelFEER (top right). On the top left, three partially methylated reads aligning to a 500 bp marker are depicted. For CelFiE, the input is given in two numbers, one equalling the sum of methylated reads at each CpG site and the other equalling the sum of the total amount of reads at each CpG site. For CelFEER, the read averages 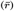 are first rounded to the closest value in {0,0.25,0.5,0.75,1}, then one-hot encoded and summed to obtain the input. The reference data is formatted in the same way. **(D)** Underlying mechanism of CelFEER for three individuals and three cell types. On the left side of the figure, the reference and input data are depicted. On the right side, the estimated methylation percentages (top) and estimated cell type proportions (bottom) are depicted. The centre part illustrates the posterior distribution of the latent variable *z*, which indicates what cell type each separate read is derived from.

A read-based approach has been adopted in multiple other tumor fraction estimation methods, such as DISMIR (12) and EpiClass (13). Even though the effectiveness of this approach has been shown for tumor fraction estimation, it has not yet been used in the related task of cell type deconvolution.

We hypothesize that read averages can increase the sensitivity of methylation-based cell type deconvolution methods. In order to evaluate the effects of using read averages without being affected by other model decisions, we decided to adapt an existing method, CelFiE (CELl Free DNA Estimation via expectation-maximization) by Caggiano et al. (8). CelFiE has the advantages that it is able to estimate missing cell types and that it can estimate cell type proportions of cfDNA with a low read coverage. Caggiano et al. demonstrated possible clinical applications of CelFiE by showing its ability to differentiate between pregnant and non-pregnant women by their proportion of placenta derived cfDNA, as well as between ALS patients and healthy individuals by their proportion of skeletal muscle cfDNA. In their work, Caggiano et al. used whole genome bisulfite sequencing (WGBS) of reference cell type DNA and input cfDNA. Since WGBS data covers the entire genome, it has the benefit that it can be used for cell type-specific biomarker discovery by comparing the methylation in all CpG sites (14).

We find that the selection of appropriate cell type markers is of crucial importance for the model performance. Using the entire genome as model input is not only computationally infeasible, but it will also likely have a negative impact on performance when CpG sites that are not informative of the cell type origin are included. By redefining the cell type informative markers, we improved CelFiE and were able to achieve better results than those reported in the original publication. The new set of markers is found using regions of 500 bp instead of single CpG sites, and only includes hypomethylated markers.

In this research, we adapted CelFiE to work at the resolution of single reads by changing the input to the average methylation value of single reads and by changing the underlying distributional assumptions accordingly. The complete workflow of the resulting method, named CelFEER (CELl Free DNA Estimation via Expectation-maximization on a Read resolution), is depicted in Figure 1a. We compared CelFEER to CelFiE on generated data and on simulated cell type mixtures composed of real WGBS data. We further applied CelFEER on the cfDNA of four ALS patients and four controls, and found that CelFEER detects a significant difference in the proportion of skeletal muscle cfDNA. Our experiments indicate that read averages can indeed more sensitively detect rare cell types. The source code for CelFEER is available at https://github.com/pi-zz-a/CelFEER

## MATERIALS AND METHODS

### CelFiE overview

As CelFEER is an adaptation of CelFiE, understanding this original method is essential for understanding CelFEER. CelFiE uses an expectation-maximization (EM) algorithm to learn the parameters of a Bayesian model of the cell type proportions of cfDNA mixtures. It does this by learning these proportions simultaneously with the average methylation percentage of each cell type at each CpG site. The methylation percentages correspond to the fraction of reads that are methylated at a specific CpG site, and are initialized by transforming the reference data into fractions. The methylation percentages are estimated because the reference cell type data is assumed to be imperfect and incomplete; CelFiE aims to learn the true methylation percentages from both the cfDNA input and the reference cell type data. The reference data consists of the methylation counts of *T* cell types indexed by *t* at *M* CpG sites indexed by *m*. More precisely, it takes the form of two *T ×M* matrices: *Y* and *D*^*Y*^, where *Y*_*tm*_ and 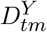 are the number of methylated and total reads at CpG site *m*, respectively, in reference cell type *t*. The reference data are assumed to be drawn from a binomial distribution for each CpG site, where the number of trials equals the reference read depth and the probabilities the true methylation percentage in the cell type of origin.

CelFiE learns the cell type proportions of multiple individuals simultaneously, allowing the method to infer information from other individuals’ methylation values. The cfDNA data from *N* individuals indexed by *n* is given in two *N ×M* matrices, *X* and *D*^*X*^. *X*_*nm*_ and 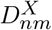 are the number of methylated and total reads at CpG site *m*, respectively, for individual *n*. An example of how these matrices are formatted is given in Figure 1c. Each *x*_*nmc*_ refers to the methylation value of a specific read *c* and can thus be either 0 or 1. These methylation values are assumed to be drawn from a Bernoulli distribution governed by the methylation percentage in the cell type of origin. 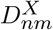 consists of the sum of all *x*_*nmc*_ while *X*_*nm*_ is the sum of all *x*_*nmc*_ that are equal to 1.

CelFiE estimates two parameters: *α* and *β*, where *α* is the final output of the model. *α*_*nt*_ is the fraction of cfDNA in person *n* that originated from cell type *t*, and *β*_*tm*_ is the true unknown methylation percentage of cell type *t* at position *m*. CelFiE models the input cfDNA as a mixture of different cell types. Whether this input originates from a given (or unknown) cell type is modeled using a latent variable *z*, where *z*_*tmc*_ = 1 when *x*_*mc*_ originates from cell type *t* and 0 otherwise. The objective is thus to describe the joint distribution *P*(*X,z,Y* |*α,β*). For the complete mathematical description of the model and its underlying assumptions, refer to the supplementary and (8).

The model iteratively relates the input to probable cell types in the expectation step, and calculates the parameters that make the input and reference data most likely in the maximization step. More mathematically put, in the expectation step the posterior distribution 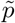 of *z* given the input data *x* and parameters *α* and *β* is calculated. These parameters are then updated by the *α* and *β* values that maximize the expectation of the joint likelihood under the calculated posterior.

### Read-based approach

CelFEER uses essentially the same model as CelFiE but with read averages as input. This changes the underlying distributions of the model, while the overall structure of the algorithm remains the same. The algorithm is visualized in Figure 1d. In CelFEER, the single counts per CpG site are replaced by five counts per 500 bp region. Each count 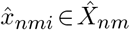 for individual *n* mapping to region *m* equals the number of reads with a discretized read average *i*, where *i*∈ {0,0.25,0.5,0.75,1}. A read average is calculated by dividing the number of methylated CpG sites by the total number of CpG sites on a read, where only reads with three or more CpG sites are used. This heuristic is adopted from previous methods (11, 12). The read average is then rounded to the closest value E.g. a read *c* from individual *n* mapping to region *m* with one out of three CpG sites methylated (and therefore a read average of 1/3) would be represented as 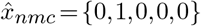. Hence each read is effectively one-hot encoded. Summing all one-hot encoded reads that fall into the same 500 bp region results in the five counts which are used as input to the model. This process is depicted in Figure 1c.

The reference data has the same composition as the input data, but instead of a set of counts per individual per site, the reference data contains a set of counts per cell type per site. Since the reference data has a different format in CelFEER compared to CelFiE, the *β* values take on a different form as well. 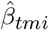 is now the proportion of reads originating from cell type *t* and mapping to region *m* that have a read average *i*.

As in CelFiE, the model aims to describe the joint distribution of the input 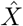, the reference *Ŷ* and the latent variable *z*, which are all assumed to be independent. In order to describe the full data likelihood, we first split it into three parts: 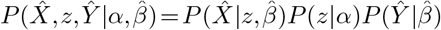.

The first part, 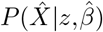, describes the likelihood of observing the read averages given that we know what cell type each read originates from and how the read averages of each cell type are distributed across the 500 bp windows. In this likelihood we look at each read *c* individually, and not yet at the total counts of all reads in a region. The probability for a read *c* at region *m* to have the value 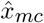 can be described as a categorical distribution where each category corresponds to a possible read average and 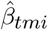 is the probability of originating from cell type *t* and belonging to category *i*. This holds for every individual *n*:

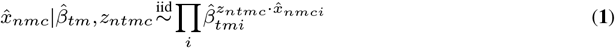

The different cell types, individuals, reads and regions are all assumed to be independent. The log-likelihood of the first part can hence be calculated as follows:

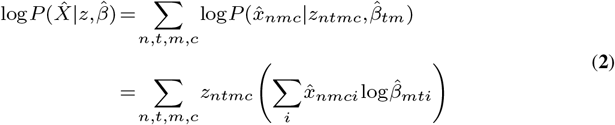

The second part of the full likelihood describes how likely it is that a read *c* originates from each cell type *t*. The probability of observing a specific cell type in the cfDNA is governed by the cell type proportions. This probability can be described using a Bernoulli distribution:

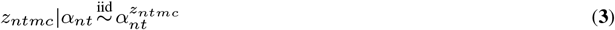

Which makes the second part of the log-likelihood:

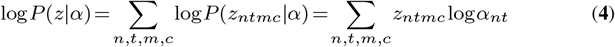

The final term is the only term that does not depend on the latent variables *z*. The reference data is assumed to be multinomially sampled with probabilities 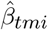 and a number of trials equal to the reference read depth, which can be obtained by summing over all read average counts:

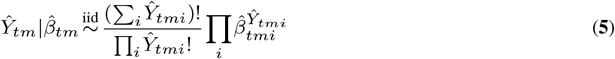

Which makes the third part of the full data likelihood equal to:

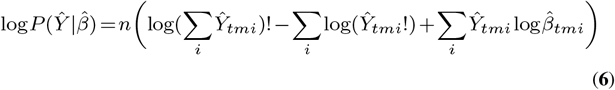

Because of the presence of the latent variables *z*, there is no closed form solution for maximizing the log-likelihood (15). Instead, we maximize the expected value of the log-likelihood under the posterior distribution of these latent variables using the EM algorithm. The posterior distribution of the latent variable *z*_*ntmc*_ is calculated by applying the Bayes rule as follows:

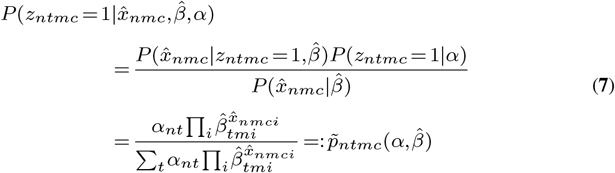

Where the distribution of 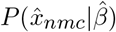 follows from the fact that each read originates from only one cell type *t*, thus summing over all cell types gives the full data distribution of the reads.

Since the read averages are one-hot encoded, there will be five possible values for the posterior 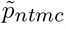. Following from this fact, we can remove the read index *c* and can start looking at the total sum of reads that have either of the five possible read averages. For each read *c* where 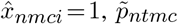 will be equal to:

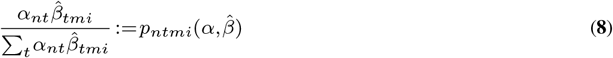

For the expectation step in the EM algorithm, we need to define the expectation of the latent variable *z* over the full data likelihood at iteration *j*.

Let *α*^(*j*)^ and *β*^(*j*)^ equal the parameters estimated at iteration *j*, and *p*^(*j*)^ := *p*(*α*^(*j*)^,*β*^(*j*)^). The expectation, also called the *Q* function, is derived in the supplementary and is defined as follows:

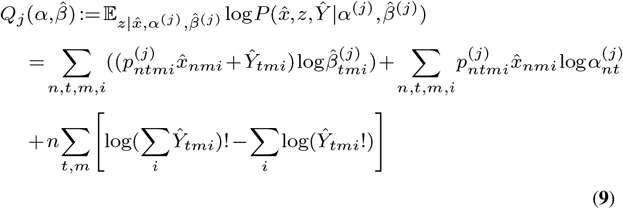

Finally *α* and 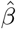 are updated by maximizing 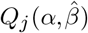, resulting in the following update formulas. For the full derivation, see the supplementary.

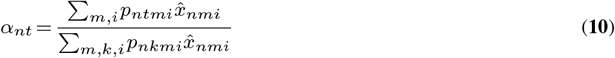

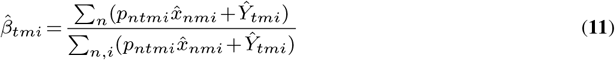

Each run of CelFEER performs the optimization 10 times independently, because EM is not guaranteed to converge to a global optimum. The log-likelihood is compared for each restart and CelFEER returns the output from the restart with the highest log-likelihood. In all simulations, we run CelFEER 50 times to capture the variance of the model output.

### Marker selection

The markers define which CpG sites will be used as input to the model. The methylation values of CpG sites at marker locations should be consistently different for different cell types, such that the methylation values at these sites can be used to distinguish between cell types. The markers are found using an adaptation of the method used by Caggiano et al. The complete process of adapting the markers is described in the supplementary. The original method (before adaptation) works as follows: All CpG sites are compared by measuring the distance between the methylation percentage of one cell type to the median methylation percentage of all cell types. The 100 markers with the largest distance are then selected as markers. Consequently, the total amount of markers found equals 100 times the number of cell types. The markers have to satisfy three requirements in the original method; the first is that a marker is only allowed to be a marker of one cell type. If the same CpG site is in the top 100 of two or more cell types, that site is not used as a marker. The second requirement is that each cell type should have at least one read at a marker location. The last requirement enforces that the median read depth of all cell types at a marker position equals at least 15.

This last requirement, however, still allows the cell type for which the CpG site is a marker to have a read depth less than 15, as long as the median read depth of all cell types is sufficient. A CpG site could be a marker for a cell type as long as it is covered by at least one read in that cell type. To remove the possibility of getting this type of marker, we introduced an extra check to ensure this cell type has a read depth at least as large as the median read depth threshold. Besides, we included one more requirement to ensure marker uniqueness. Instead of comparing only the top 100 markers of each cell type, we compared the top 150 markers of each cell type. After this comparison, again only the top 100 markers are used. This extra step prevents the situation where a marker is in the top 101 of one cell type and in the top 99 of another, which could lead to the inclusion of less differential markers.

The original method should, in theory, be able to find both hypo- and hypermethylated markers. In practice, it finds almost solely hypomethylated markers. Comparing each cell type’s methylation percentage to the median methylation percentage can make markers less distinct, as is shown in Figure S1c. Therefore, we adapted the method to compare each cell type’s methylation percentage to the minimum methylation percentage of all other cell types, as is shown in Figure S1d. We found that hypomethylated markers are best at differentiating between cell types (see supplementary).

Originally, CelFiE uses as input and as reference data the methylation values at the marker CpG sites summed with the methylation values of CpG sites in the *±*250 bp surrounding the marker sites. We improved CelFiE by first summing the CpG sites into 500 bp windows which are subsequently used to find marker regions. Otherwise, markers on regions are found using the exact same approach as markers found on single CpG sites. The difference between finding markers on single CpG sites and on regions is shown in Figures S1a and S1b. As there is no requirement for the amount of CpG sites in a region and only for the minimum read coverage of a region, the amount of CpG sites per marker can differ. Because summing the CpG sites into 500 bp windows substantially increased the read coverage at potential marker regions, we increased the read depth threshold to 150. To find the value for this threshold, we tried a range of increasing values and compared the resulting markers by their distance between cell types.

Finding the markers using the read average data largely follows the same approach. First, the chromosome is split into 500 bp windows into which the reads are mapped. For each cell type, the read averages are averaged over all reads that map to the same window. The CelFEER markers are found by comparing these averages. This process is illustrated in Figure 2. For the read averages, we again optimized the read depth threshold and observed that the best markers were found using a read depth threshold of 20. The large difference with the read depth threshold for the CelFiE input (after summing in 500 bp windows) can be explained by the large difference in the scale of the input of CelFiE and CelFEER. This difference in scale is due to the fact that all CpG sites on a read contribute to a single value in CelFEER, and to multiple values in CelFiE.

**Figure 2.**
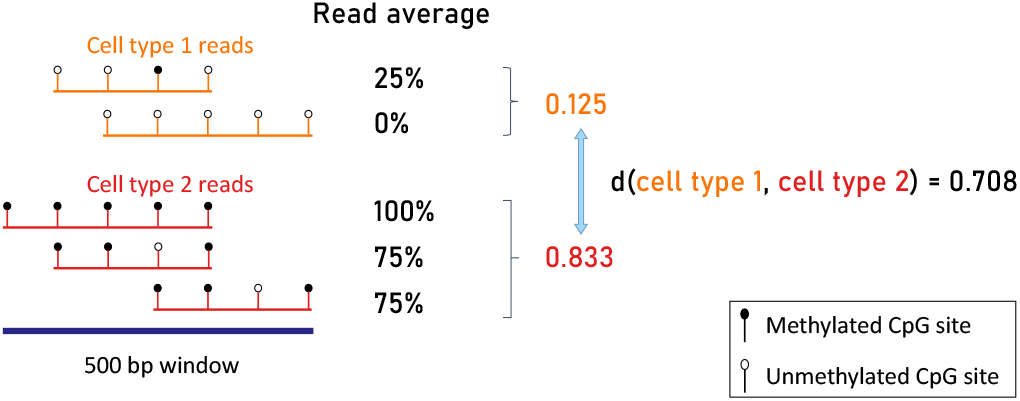
Illustration of the method for determining the distance between two cell types using read averages. First, the average of all read averages is determined for each cell type. These are then compared to find the distance between cell types.

**Figure 3.**
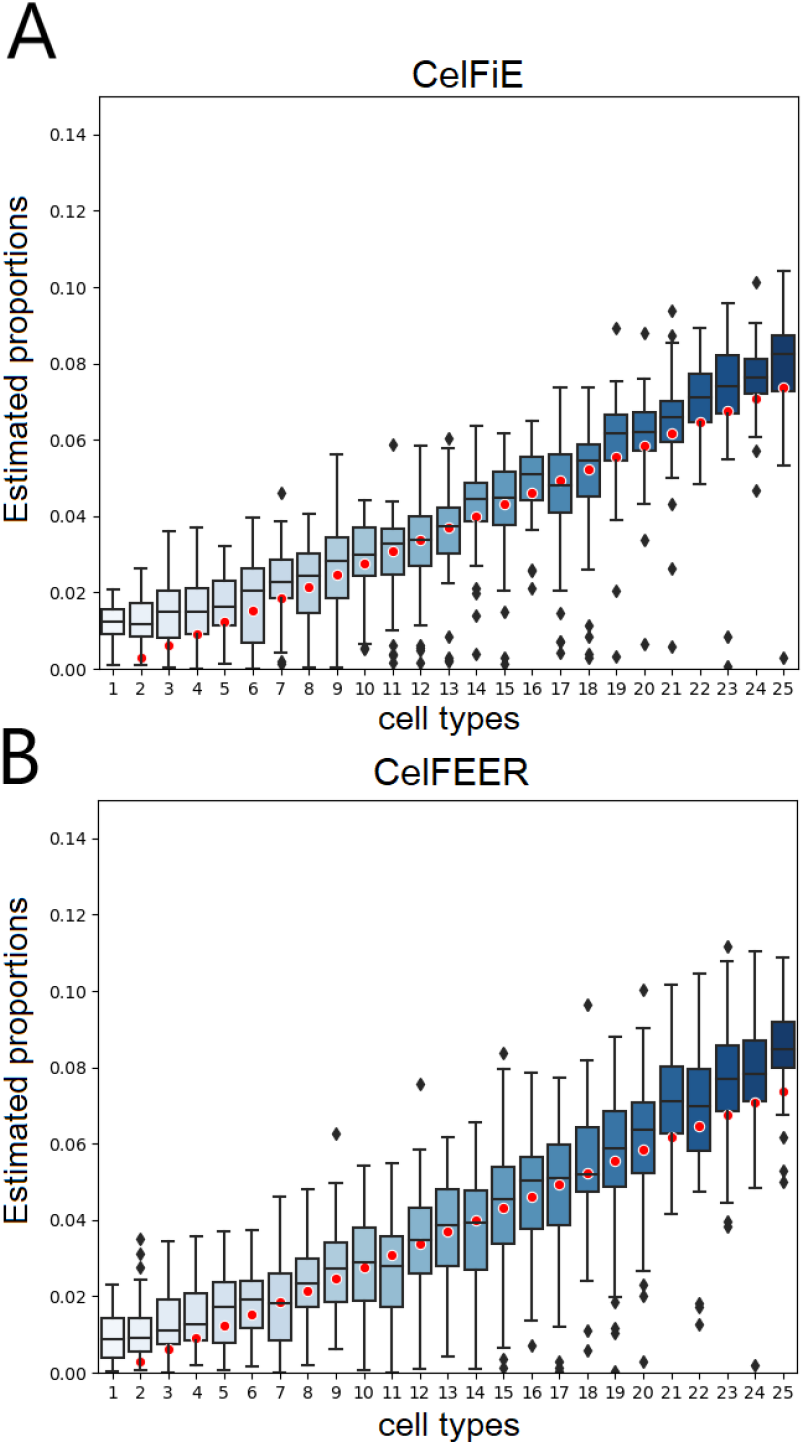
Simulations on generated data for one individual. Each boxplot displays the estimated proportion of a cell type for replicate model runs. The red dots indicate the true cell type proportions for 25 cell types.

Since the approach to summarize read averages into bins is slightly different from the approach used to bin the CpG count data, we bin the CpG count data in the same manner as the read averages when comparing CelFiE and CelFEER.

### Simulations with artificial data

In order to validate if CelFEER works under the model assumptions, simulations with artificial data were set up as follows.

The input and reference data are generated according to the distributions assumed by the model. The simulations use the same parameters as originally used by Caggiano et al. in their artificial simulations. In each random restart, *α* is randomly initialized by drawing from a uniform distribution and normalizing to ensure the values sum to one. *β*_*i*_ is initialized by taking 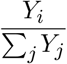.

We also ran simulations where one or two cell types are considered unknown, i.e. they are not included in the reference. In these cases, we created the true cell type proportions and true cell type methylation as before. In the reference data that is passed to CelFEER, the methylation values for the unknown cell type are set to {0, 0, 0, 0, 0}. This way, the estimated methylation percentages for an unknown cell type are initialized to {0.2, 0.2, 0.2, 0.2, 0.2}.

### Simulations on WGBS data

To further evaluate the method, we simulated cfDNA data by mixing WGBS data of different cell types. The cell type data was obtained from ENCODE (16) and Blueprint (17), and is composed of T-cell CD4, monocyte, macrophage, memory B cell, neutrophil, adipose, pancreas, small intestine, stomach and tibial nerve data. The sample identifiers of the used data can be found in Table S1. The data is a mixture of paired-end and single-end reads, and consists of the same datasets used by Caggiano et al. For each cell type, one sample was used to compose the reference matrix and one to simulate a cfDNA mixture. Both sex chromosomes were removed, to make the reference matrix applicable to both sexes and to ensure that random methylation due to X chromosome inactivation is not seen as relevant. Furthermore, all SNPs in dbSNP (18) were removed.

To ensure that each dataset contained an equal amount of reads before creating a mixture, the total read coverage of each cell type was normalized by dividing by the total amount of reads of all cell types and multiplying with the average amount of reads. Next, the methylation values of each cell type were multiplied with the desired proportion for that cell type. These proportions were always ensured to add up to one by dividing each cell type’s proportion by the sum of all cell types’ proportions.

In the original publication (8), WGBS mixtures were created in a similar manner, with the difference that we corrected for differences in read depth among the different cell types before downsampling the reads.

The mixtures of read averages were created similarly. First, all read counts were normalized such that each cell type occurred in equal quantities before multiplying the input with the desired proportions.

For both methods the reference data was not normalized. During parameter convergence, the only equation where the reference data is used is Equation **11**, where it is transformed to a proportion. The absolute counts of the reference data only matter in their proportion to the input data in Equation **11**. It does, however, make sense to not normalize the reference data here since it is logical that reference data with a higher coverage is more reliable and should therefore weigh more in the calculation of *β*.

## RESULTS

### Simulations using generated data

To test whether CelFEER works as expected, we followed Caggiano et al. in generating data to simulate cfDNA input and cell type DNA reference data. Using generated data, they showed that CelFiE (i) estimates proportions correlated to the true cell type proportions, (ii) is able to detect small differences between two groups of individuals and (iii) is able to estimate the proportions of unknown cell types (i.e. cell types that are present in the input data, but not in the reference).

The results of these simulations are not an accurate reflection of the model performance, as the simulations for neither CelFiE nor CelFEER model any correlation between sites. As a result, the input of adjacent sites is not summed together as is done for WGBS data, even though Caggiano et al. have shown that the original method does not return sensible results on WGBS data without summing adjacent sites. The simulations do serve as a way of investigating whether CelFEER has the same three properties (which are described above) as CelFiE.

#### CelFEER estimates of generated data correlate to true proportions

As a first evaluation of the read-based method, the performance of CelFEER is compared to the performance of CelFiE on generated data. The simulations follow the approach of (8), meaning that 50 replicates were run, each with 25 cell types, 6000 CpG sites and 1 individual. The read depth at each CpG site was drawn from a Poisson distribution centred around 10.

To compare the performance, we measured the Pearson’s correlation between the estimated and true cell type proportions. CelFEER performed slightly worse, with a mean Pearson’s correlation *r*^2^ = 0.84*±*0.05 compared to *r*^2^= 0.87*±*0.07 for CelFiE. The result of CelFiE found by us is, however, not as good as the result reported in (8), where the supposedly same simulations result in *r*^2^ = 0.96*±*0.01.

#### CelFEER and CelFiE do not detect a significant difference between two groups

Even in individuals with cfDNA originating from aberrant cell types, most of the cfDNA is usually derived from hematopoietic origins (10). In other words, the actual amount of cfDNA from an aberrant cell type can be very small. Therefore, it is important to be able to differentiate between a group that does not have this cell type and a group that has only a very small amount of it. To this end, we simulated a cell type that made up a proportion of 0.01 of the cfDNA of five individuals (group A) and 0 of the cfDNA of five other individuals (group B). Ten cell types were used in total on an input of 1000 CpG sites. The remaining nine cell types had a true proportion drawn from a uniform distribution between 0.5 and 1, which were then normalized such that all proportions summed to one.

Figure S2 shows the estimated proportion of the rare cell type for both groups, using both CelFiE and CelFEER. Averaged over 50 replicates, CelFiE estimated a proportion of 0.03*±*0.01 in group A and 0.025*±*0.007 in group B, while CelFEER estimated proportions of 0.031*±*0.01 and 0.026*±*0.008 for the two groups respectively. A two-samples t-test done for each individual showed no significant difference between the average proportions estimated by both methods in neither groups (*p*> 0.1 for all individuals). Moreover, the proportions of the rare cell type are highly overestimated in both groups.

#### CelFEER estimates proportions of unknown cell types

One of the advantages of CelFiE over previous deconvolution methods is its ability to infer cell type information from the methylation states of other individuals. This way it can estimate the cell type proportions of cell types that are not present in the reference data. As in the original paper, we generated cfDNA for 1000 CpG sites, 10 cell types and 10 individuals at a read depth of 10. In the reference data, we set the methylation states of the last cell type to 0 at each CpG site. The true proportion of this unknown cell type was drawn from a normal distribution centred around 0.2 with a standard deviation of 0.1, and clipped if smaller than 0 or larger than 1. All other cell type proportions were drawn from a uniform distribution between 0 and 1, and together with the unknown cell type the proportions were made to sum to 1. This was done for each individual separately.

We measured the root mean squared error (RMSE) of the estimated proportion of the missing cell type. Averaged over all individuals, CelFEER resulted in an RMSE of 0.0009, and CelFiE in an RMSE of 0.0010. This shows that CelFEER is also capable of estimating proportions of unknown cell types in generated data.

### Results of simulations using WGBS data

Since there are no ground truth cell type proportions available for real cfDNA data, we simulated mixtures of cfDNA by mixing WGBS data of different cell types at known proportions. To this end, we used the same data used by Caggiano et al. However, we were limited to using seven different cell types because of the availability of read data at the time of testing.

#### Comparison between CelFiE and CelFEER

To compare the performance of CelFEER to the performance of CelFiE, we again simulated cfDNA mixtures by artificially mixing WGBS data of different cell types. Again we followed the same approach as CelFiE to create the true cell type proportions for 100 individuals. The marker regions of both models were found using their reference data and were therefore different for the two models, since one set of regions was found by comparing CpG site averages and the other by comparing read averages of different cell types.

In Figure S3, the results of 50 replicate runs for a randomly selected individual are shown. Without unknown cell types in the reference data, CelFEER results in a correlation of *r*^2^ = 0.94*±*0.04 while CelFiE results in a correlation of *r*^2^*±* 0.86*±*0.09, which is higher than the performance reported in the original CelFie publication (8). We find that the difference in correlation between CelFEER and CelFiE is significant; *t*(9998) = 58.11,*p*< 0.001. To examine whether this would go at the expense of runtime, we measured the time it takes each method to run one replicate. On our system, CelFEER requires ∼1.1 times the time needed by CelFiE.

Since one of the assets of CelFiE is its ability to infer the proportions of unknown cell types, we expected CelFEER to outperform CelFiE on this aspect as well. Similar to the original experiments in (8), we masked T cells in the reference data by setting all T cell reference methylation values to 0. CelFEER highly overestimates the missing cell type proportion and therefore estimates proportions that are less correlated to the true cell type proportions than CelFiE does, although CelFiE also overestimates considerably (see Table 1 and Figure S3). When small intestine cells are masked as well, the correlation between the estimated and true cell type proportions decreases even more.

**Table 1.** Pearson’s correlation (*r*^2^) between true and estimated cell type proportions (*α* estimates) and between reference methylation and estimated methylation values (*β* estimates) of a simulated mixture of seven different cell types.

| Unknowns | Parameter | CelFiE $r^2$ | CelFEER $r^2$ |
| --- | --- | --- | --- |
| 0 | $\alpha$ | $0.86 \pm 0.09$ | $0.94 \pm 0.04$ |
| | $\beta$ | $9.98e-1 \pm 0.03e-1$ | $0.93 \pm 0.03$ |
| 1 | $\alpha$ | $0.60 \pm 0.19$ | $0.48 \pm 0.25$ |
| | $\beta$ | $0.92 \pm 0$ | $0.89 \pm 0.11$ |
| 2 | $\alpha$ | $0.30 \pm 0.34$ | $0.19 \pm 0.29$ |
| | $\beta$ | $0.85 \pm 0.25$ | $0.77 \pm 0.26$ |

In addition to comparing the estimated cell type proportions and their correlation to the true proportions, we investigated the estimated cell type methylation values. We measured the correlation between the estimated cell type methylation percentages and the methylation percentages obtained by normalizing the methylation values of the reference data to sum to one. It is remarkable how this correlation is consistently higher for CelFiE (Table 1). This implies that the methylation percentages estimated by CelFiE diverge only very little from the reference methylation. This probably means that CelFEER takes the input of other individuals more into account when estimating the methylation values, and therefore indirectly when estimating the cell type proportions.

Another advantage of CelFiE over previous methods is that it works with low coverage input data. A higher read coverage means higher sequencing costs, and it is therefore desirable that CelFEER performs sufficiently on low coverage data as well. To test this, before mixing the cell types we normalized the read coverage of each cell type to equal the total amount of input regions multiplied with a constant, *n*. This way, each cell type covered each region with *n* reads on average. For each *n*∈ {2,5,10,50} the average correlation over 50 replicates and 100 individuals was measured. The cell type proportions were generated in the same manner as before, and no unknowns were estimated. The relation between the correlation and the coverage is shown in Figure 4. We can conclude that for a stable performance, the coverage should be 10 or higher. Interestingly, the correlation between the estimated and true cell type proportions increases a little for CelFiE when *n* = 5. It is possible that lowering the coverage acts as a noise reduction on the CelFiE input. Even on the lowest coverage, CelFEER outperforms CelFiE, showing that CelFEER is a suitable method for low coverage data.

**Figure 4.**
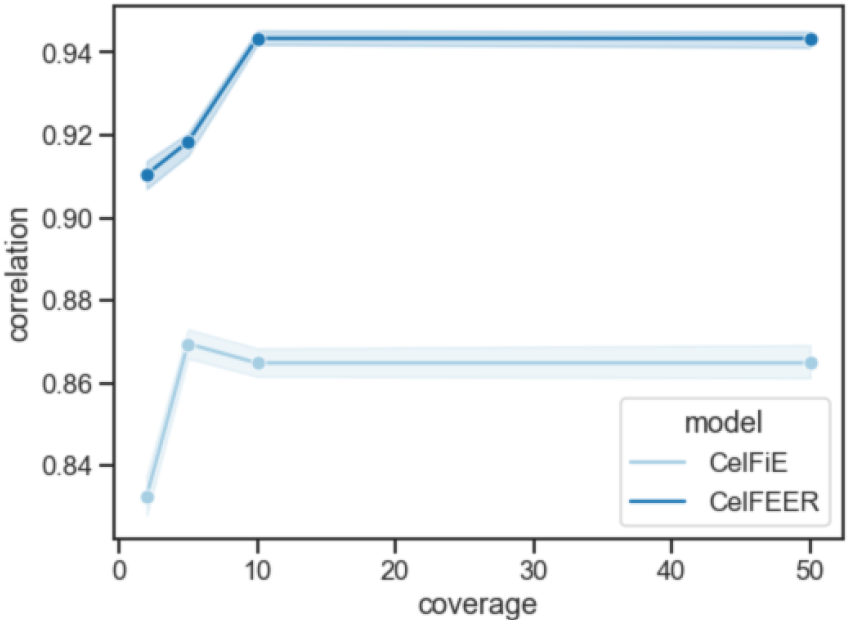
Relation between the input coverage and the correlation between the estimated and true cell type proportions. The full range of the correlations of 100 individuals and 50 replicates is highlighted.

#### Markers found on read averages are different from markers found on count input

Finally we were interested in comparing the markers found using read averages to the markers found using CpG site averages. We hypothesised that CelFEER works better with markers found on the read averages of the reference data, on the grounds that CelFEER differentiates cell types by their read averages. Additionally, as reasoned in the introduction, read averages should be more sensitive to differences in methylation status between cell types. We again performed the same experiments, using a simulated mixture of seven different cell types.

We firstly checked the overlap in markers found using both methods. Of the 700 markers, 130 markers were found by both. Each of the seven cell types has markers that are found by both methods. There are no regions that are a marker for one cell type in one method and a marker for another cell type in the other method.

Using the markers found by CelFiE, CelFEER performed similarly with a correlation of *r*^2^ = 0.94*±*0.04 (Figure S4). The correlation between the cell type proportions estimated by CelFiE using CelFEER’s markers is *r*^2^ = 0.69*±*0.21, indicating that the markers found by CelFEER are not suitable for the input of CelFiE. Averaged over all cell types, the difference in methylation percentage between cell types at CelFiE’s marker locations is 0.65 for both the reference and input data, where the reference data showed slightly less variation with a standard deviation of 0.19 compared to 0.20 for the input data. For CelFEER, this difference is 0.66*±*0.20 for the input and 0.64*±*0.22 for the reference. Figure S5 does show that for some cell types the variation in the distance from the median is substantially larger for the CelFEER markers.

### Application in ALS

Caggiano et al. showed that CelFiE is able to differentiate between Amyotrophic Lateral Sclerosis (ALS) patients and a control group by the estimated proportion of skeletal muscle derived cfDNA. Although it is interesting to see if CelFEER is also able to distinguish between the ALS and the control group, it is hard to evaluate the method based on its cell type proportion estimates since there are no ground truth cell type proportions available. Moreover, while Caggiano et al. used 28 case and 25 control samples, we only used four case and four control samples. The reference data consists of all 19 cell types given in supplementary Table S1.

We firstly fully decomposed the cfDNA, thus estimating the proportions of each of the 19 cell types present in the reference. The five cell types with the highest proportions estimated by CelFEER were, in both groups, the following: neutrophil, monocyte, erythroblast, spleen and eosinophil. CelFiE estimated similar proportions, but instead of spleen it estimated adipose to be the fourth highest in proportion. In their own work (8), however, neither spleen nor adipose, but macrophage cells are in this top five. Still, these results mostly correspond to the findings of Moss et al. (10). The full decomposition can be seen in Figure 5a and Figure 5b.

**Figure 5.**
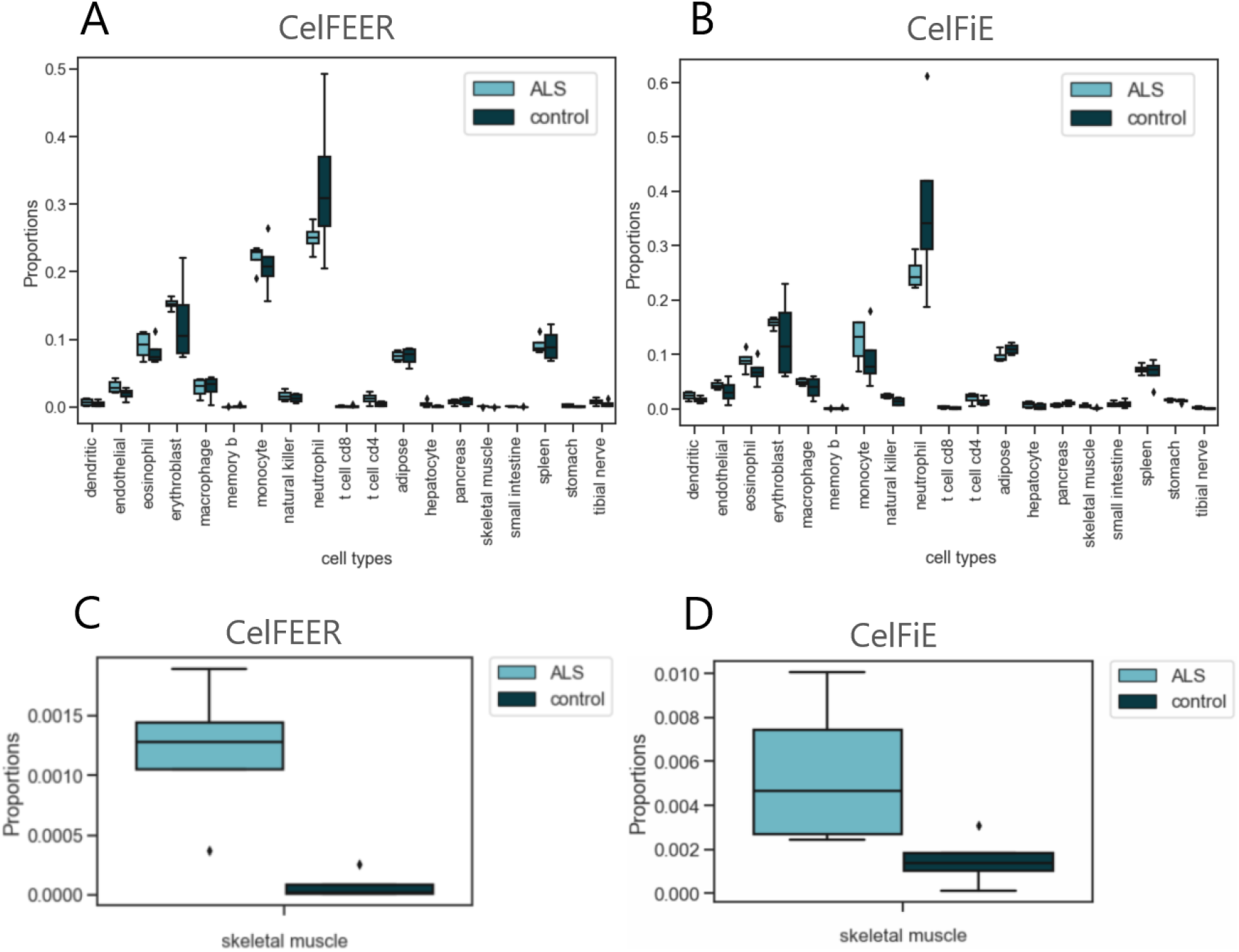
Estimated proportions of cfDNA in ALS patients (*n* = 4) and a control group (*n* = 4). **Figures A, B** The complete cell type decomposition of CelFEER and CelFiE respectively. **Figures C, D** The estimated proportion of skeletal muscle cfDNA by CelFEER and CelFiE.

Next, we specifically examined the skeletal muscle cell proportions in both groups. CelFiE estimated an average proportion of 5.5e −3*±*3.1e− 3 in the ALS case group, and 1.5e− 3*±*1.1e− 3 in the control group (Figure 5d). A two-sample t-test did not indicate a significant difference between the two groups; *t*(6) = 2.09,*p* = 0.08. CelFEER, on the contrary, did find a significant difference, with an average proportion of 1.2e− 3*±*5.4e− 4 for the ALS case group and 7.7e− 5*±*1e− 4 for the control group (Figure 5c); *t*(6) = 3.54,*p* = 0.01. Clearly, CelFEER is able to detect small fractions of rare cell types in cfDNA.

## DISCUSSION

The analysis of cfDNA has some attractive properties, such as the possibility to detect and monitor disease without undertaking invasive biopsies (2). By retrieving the cell types of origin of cfDNA, it is possible to obtain a complete overview of all cells that shed cfDNA, and even of the amount of cfDNA each cell type yields. An inquiry in the cell type proportions can indicate the presence of aberrant cell types, such as tumor cells, in the cfDNA. Yet, detection of aberrant cell types can be difficult, especially in early stages of disease. Recent methods use the methylation states at CpG sites that cause a differential gene expression in different cell types. In this research, we adapted one such method, CelFiE (8), to instead use differential methylation averages of individual reads. The intuition behind this approach is that the methylation averages of individual reads differentiate more than CpG site averages, since aberrant reads are almost undetectable when averaged with healthy reads. This new method, named CelFEER, uses an expectation-maximization algorithm and a reference cell type dataset to estimate the true cell type proportions of cfDNA mixtures. We showed that CelFEER performs as expected on generated data, and outperforms CelFiE on cfDNA simulated using mixtures of WGBS data. Moreover, it can accurately differentiate between ALS patients and a control group by comparing the proportions of skeletal muscle derived cfDNA in the two groups.

The model’s performance is highly reliant on the quality of the input regions, where the quality is defined by the difference in methylation between cell types at an input region. In pursuit of improving CelFiE’s model performance, we improved the original method for finding markers by applying the following changes: (i) we differentiated between 500 bp regions instead of single CpG sites, (ii) we focused on hypomethylated regions and (iii) we applied stricter rules to marker regions. To find marker regions for CelFEER, we devised a method that largely follows the same approach as CelFiE but instead uses the read averages of the reference data.

The read averages are formulated in a way that one read average, so one single value, summarizes multiple CpG sites. For this reason, the range of the input is much lower for CelFEER than for CelFiE. In addition, CelFEER filters out reads covering less than three CpG sites, which decreases the range even more. It may be interesting to investigate whether allowing for reads with a lower CpG site coverage gives improvements to the model. Low read quality is one of the disadvantages of working with WGBS data, as the bisulfite conversion is known to be detrimental to the DNA (19). Another way for compensating for the smaller range would be to increase the amount of samples used in the reference dataset. Currently, each reference cell type consists of the DNA of a single individual.

If the reference data does not include all of the cell types found in the cfDNA, the proportions of the cell types that are included will be overestimated. Since actual cfDNA is likely to contain a component of cell types that are absent from the reference data (8), it is useful to estimate proportions of unknown cell types. However, CelFEER currently greatly overestimates the proportions of unknown cell types. It may be possible to improve this by changing the input for unknown cell types, as we presently employ CelFiE’s method of setting unknown cell types to 0, which may not work for CelFEER. In relation to that, we may need to change the initial values for the estimated methylation percentages for unknown cell types. Despite the improvements made to the selected marker regions, there is potential for more distinct markers, perhaps by adopting a completely new approach. After all, the method for finding markers was optimized for CpG count data and then translated almost exactly to read average data. Read averages may, however, require a completely different approach for finding markers, such as the switching reads defined by Li et al. (12). An adequate set of differential regions not only improves model performance but also allows for targeted sequencing of these regions only, for example using RRBS, and can thus reduce the sequencing cost (20).

We chose to discretize the read averages into five bins instead of treating them as continuous values. This substantially speeds up the method, because it means that we only need to estimate the distribution over five possible read averages instead of all possible read averages. Moreover, binning ensures we have more evidence for each of the five distributions to be estimated. Although the input size of CelFEER is larger than the input size of CelFiE (read averages are described by five counts instead of the two counts used by CelFiE), it suffers only from a minor increase in runtime. Like CelFiE, CelFEER is an efficient method that scales linearly in the size of the input and reference. Even so, it could be beneficial to consider CelFEER’s performance when using more or less counts. Using less counts, i.e. rounding the read averages more before summing similar averages, would likely decrease model performance but speed up computations. Using more counts, on the other hand, may give an increase in performance that is worth the added computation time.

Finally, the use of CelFEER in practical applications should be investigated further by testing the model on more cfDNA data. A first step would be to use more samples in the ALS experiment. Eventually the model could be tested on, for instance, pregnancy and cancer samples.

In conclusion, with CelFEER, we showed that a cell type deconvolution method can more sensitively estimate cell type proportions when using read averages instead of CpG site averages, even at a low input read coverage.

## Conflict of interest statement

None declared.

## Notes

### Competing Interest Statement

The authors have declared no competing interest.

## REFERENCES

1. Snyder, M. W., Kircher, M., Hill, A. J., Daza, R. M., and Shendure, J. (2016) Cell-free DNA comprises an in vivo nucleosome footprint that informs its tissues-of-origin. Cell, 164(1-2), 57–68.

2. Lo, Y. M. D., Han, D. S. C., Jiang, P., and Chiu, R. W. K. (2021) Epigenetics, fragmentomics, and topology of cell-free DNA in liquid biopsies. Science, 372(6538).

3. Sun, K., Jiang, P., Chan, K. A., Wong, J., Cheng, Y. K., Liang, R. H., Chan, W.-k., Ma, E. S., Chan, S. L., Cheng, S. H., et al. (2015) Plasma DNA tissue mapping by genome-wide methylation sequencing for noninvasive prenatal, cancer, and transplantation assessments. Proceedings of the National Academy of Sciences, 112(40), E5503– E5512.

4. Barefoot, M. E., Loyfer, N., Kiliti, A. J., McDeed IV, A. P., Kaplan, T., and Wellstein, A. (2021) Detection of Cell Types Contributing to Cancer From Circulating, Cell-Free Methylated DNA. Frontiers in genetics, 12.

5. Li, B., Pei, G., Yao, J., Ding, Q., Jia, P., and Zhao, Z. (2021) Cell-type deconvolution analysis identifies cancer-associated myofibroblast component as a poor prognostic factor in multiple cancer types. Oncogene, 40(28), 4686–4694.

6. Greenberg, M. V. and Bourc’his, D. (2019) The diverse roles of DNA methylation in mammalian development and disease. Nature reviews Molecular cell biology, 20(10), 590–607.

7. Shoemaker, R., Deng, J., Wang, W., and Zhang, K. (2010) Allele-specific methylation is prevalent and is contributed by CpG-SNPs in the human genome. Genome research, 20(7), 883–889.

8. Caggiano, C., Celona, B., Garton, F., Mefford, J., Black, B. L., Henderson, R., Lomen-Hoerth, C., Dahl, A., and Zaitlen, N. (2021) Comprehensive cell type decomposition of circulating cell-free DNA with CelFiE. Nature communications, 12(1), 1–13.

9. Kang, S., Li, Q., Chen, Q., Zhou, Y., Park, S., Lee, G., Grimes, B., Krysan, K., Yu, M., Wang, W., et al. (2017) CancerLocator: non-invasive cancer diagnosis and tissue-of-origin prediction using methylation profiles of cell-free DNA. Genome biology, 18(1), 1–12.

10. Moss, J., Magenheim, J., and Neiman, D. e. a. Comprehensive human cell-type methylation atlas reveals origins of circulating cell-free DNA in health and disease. (Nov, 2018).

11. Li, W., Li, Q., Kang, S., Same, M., Zhou, Y., Sun, C., Liu, C.-C., Matsuoka, L., Sher, L., Wong, W. H., et al. (2018) CancerDetector: ultrasensitive and non-invasive cancer detection at the resolution of individual reads using cell-free DNA methylation sequencing data. Nucleic acids research, 46(15), e89–e89.

12. Li, J., Wei, L., Zhang, X., Zhang, W., Wang, H., Zhong, B., Xie, Z., Lv, H., and Wang, X. (2021) DISMIR: Deep learning-based noninvasive cancer detection by integrating DNA sequence and methylation information of individual cell-free DNA reads. Briefings in bioinformatics, 22(6), bbab250.

13. Miller, B. F., Pisanic II, T. R., Margolin, G., Petrykowska, H. M., Athamanolap, P., Goncearenco, A., Osei-Tutu, A., Annunziata, C. M., Wang, T.-H., and Elnitski, L. (2020) Leveraging locus-specific epigenetic heterogeneity to improve the performance of blood-based DNA methylation biomarkers. Clinical epigenetics, 12(1), 1–19.

14. Loyfer, N., Magenheim, J., Peretz, A., Cann, G., Bredno, J., Klochendler, A., Fox-Fisher, I., Shabi-Porat, S., Hecht, M., Pelet, T., et al. (2022) A human DNA methylation atlas reveals principles of cell type-specific methylation and identifies thousands of cell type-specific regulatory elements. Biorxiv,.

15. Bishop, C. M. and Nasrabadi, N. M. (2006) Pattern recognition and machine learning, Vol. 4, Springer,.

16. Consortium, E. P. et al. (2012) An integrated encyclopedia of DNA elements in the human genome. Nature, 489(7414), 57.

17. Fernández, J. M., de la Torre, V., Richardson, D., Royo, R., Puiggròs, M., Moncunill, V., Fragkogianni, S., Clarke, L., Flicek, P., Rico, D., et al. (2016) The BLUEPRINT data analysis portal. Cell systems, 3(5), 491– 495.

18. Kitts, A. and Sherry, S. (2002) The single nucleotide polymorphism database (dbSNP) of nucleotide sequence variation. The NCBI handbook. McEntyre J, Ostell J, eds. Bethesda, MD: US national center for biotechnology information,.

19. Kurdyukov, S. and Bullock, M. (2016) DNA methylation analysis: choosing the right method. Biology, 5(1), 3.

20. Beck, D., Ben Maamar, M., and Skinner, M. K. (2022) Genome-wide CpG density and DNA methylation analysis method (MeDIP, RRBS, and WGBS) comparisons. Epigenetics, 17(5), 518–530.

