## Supplement for "Cell type deconvolution of methylated cell-free DNA at the resolution of individual reads"

### Supplementary

#### 1 Equations

##### Original CelFiE equations

Posterior distribution:

$$\begin{aligned} p_{ntm1}(\alpha,\beta) &:= p_{ntmc}(\alpha,\beta) \qquad \text{if } x_{nmc}=1 \\ &= \frac{\beta_{tm}\alpha_{nt}}{\sum_k \beta_{kt}\alpha_{nk}} \\ p_{ntm0}(\alpha,\beta) &:= p_{ntmc}(\alpha,\beta) \qquad \text{if } x_{nmc}=0 \\ &= \frac{(1-\beta_{tm})\alpha_{nt}}{\sum_k (1-\beta_{kt})\alpha_{nk}} \end{aligned} \tag{1}$$

$\alpha$ and $\beta$ update formula:

$$\alpha_{nt} = \frac{\sum_m \left(x_{nm}p_{ntm1} + (D^X_{nm} - x_{nm})p_{ntm0}\right)}{\sum_{km}\left(x_{nm}p_{nkm1} + (D^X_{nm} - x_{nm})p_{nkm0}\right)} \tag{2}$$

$$\beta_{tm} = \frac{\sum_n p_{ntm1}X_{nm} + nY_{tm}}{\sum_n p_{ntm0}(D^X_{nm} - X_{nm}) + nD^Y_{tm} + \sum_n p_{ntm1}X_{nm}} \tag{3}$$

Log-likelihood formulation:

$$\begin{aligned} Q(\alpha,\beta) = &\sum_{n,t,m}\left[(Y_{tm} + p_{ntm1}X_{nm})\log(\beta_{tm}) + \left(D^Y_{tm} - Y_{tm} + p_{ntm0}(D^X_{nm} - X_{nm})\right)\log(1-\beta_{tm})\right] \\ &+ \sum_{n,t,m}\left(X_{nm}p_{ntm1} + (D^X_{nm} - X_{nm})p_{ntm0}\right)\log\alpha_{nt} \end{aligned} \tag{4}$$

##### Derivation of full data log-likelihood

$$
\begin{aligned}
Q(\alpha,\hat{\beta}) &:= \mathbb{E}_{z|\hat{X},\alpha,\hat{\beta}} \log P(\hat{X},z,Y|\alpha,\beta) \\
&= \mathbb{E}_{z|\hat{X},\alpha,\hat{\beta}} \left(\log P(\hat{X}|z,\hat{\beta}) + \log P(z|\alpha) + \log P(\hat{Y}|\hat{\beta})\right) \\
&= \sum_{n,t,m,c} \mathbb{E}_{z|\hat{X},\alpha,\hat{\beta}} \left[ z_{ntmc} \sum_i \hat{x}_{nmci} \log \hat{\beta}_{tmi} + z_{ntmc} \log \alpha_{nt} \right] \\
&+ \sum_{n,t,m} \left( \log(\sum_i \hat{Y}_{tmi}!) - \sum_i \log(\hat{Y}_{tmi}!) + \sum_i \hat{Y}_{tmi} \log \hat{\beta}_{tmi} \right) \\
&= \sum_{n,t,m,c} \tilde{p}_{ntmc} \left[ \sum_i \hat{x}_{nmci} \log \hat{\beta}_{tmi} + \log \alpha_{nt} \right] \\
&+ \sum_{n,t,m} \left( \log(\sum_i \hat{Y}_{tmi}!) - \sum_i \log(\hat{Y}_{tmi}!) + \sum_i \hat{Y}_{tmi} \log \hat{\beta}_{tmi} \right) \\
&= \sum_{n,t,m} \left[ \sum_i p_{ntmi} \hat{x}_{nmi} \log \hat{\beta}_{tmi} + \sum_i p_{ntmi} \hat{x}_{nmi} \log \alpha_{nt} \right] \\
&+ \sum_{n,t,m} \left[ \log(\sum_i \hat{Y}_{tmi}!) - \sum_i \log(\hat{Y}_{tmi}!) + \sum_i \hat{Y}_{tmi} \log \hat{\beta}_{tmi} \right] \\
&= \sum_{n,t,m,i} ((p_{ntmi}\hat{x}_{nmi} + \hat{Y}_{tmi}) \log \hat{\beta}_{tmi}) + \sum_{n,t,m,i} p_{ntmi}\hat{x}_{nmi} \log \alpha_{nt} + n \sum_{t,m} \left[ \log(\sum_i \hat{Y}_{tmi})! - \sum_i \log(\hat{Y}_{tmi}!) \right]
\end{aligned} \tag{5}
$$

##### Derivation of $\alpha$ and $\hat{\beta}$ update formulas

Maximization of the log-likelihood w.r.t $\alpha$ and $\hat{\beta}$ can be done using the following fact that for a probability simplex $S_K \subset \mathbb{R}^K$ and any $a \in \mathbb{R}^K_{++}$:

$$
\arg\max_{p \in S_K} \sum_k a_k \log p_k = (a_1, ..., a_K) / \sum_{k=1}^K a_k
$$

To derive $\alpha_t$, we let $a_t = \sum_i p_{tmi}\hat{x}_i$ s.t.

$$
\alpha_{nt} = \frac{\sum_{m,i} p_{ntmi}\hat{x}_{nmi}}{\sum_{m,t,i} p_{ntmi}\hat{x}_{nmi}} \tag{6}
$$

For $\hat{\beta}_{tmi}$ we let $a_i = p_{tmi}\hat{x}_i + \hat{Y}_{tmi}$ s.t.

$$
\hat{\beta}_{tmi} = \frac{\sum_n (p_{ntmi}\hat{x}_{nmi} + \hat{Y}_{tmi})}{\sum_{n,i} (p_{ntmi}\hat{x}_{nmi} + \hat{Y}_{tmi})} \tag{7}
$$

#### 2 Data

Tab. S1: WGBS cell type data and sources

| Cell type | Database | Sample 1 | Sample 2 |
|---|---|---|---|
| CD4-positive, alpha-beta T cell | Blueprint | S007G7 | S007DD |
| CD8-positive, alpha-beta T cell | Blueprint | C003VO | C00256 |
| endothelial cell of umbilical vein (resting) | Blueprint | S00DCS | S00BJM |
| monocyte | Blueprint | S01MAPA1 | S01E03A1 |
| erythroblast | Blueprint | S002S3 | S002R5 |
| macrophage | Blueprint | S0022I | S00390 |
| mature eosinophil | Blueprint | S00V65 | S006XE |
| memory B cell | Blueprint | C003N3 | S017RE51 |
| cytotoxic CD56-dim natural killer cell | Blueprint | C006G5 | C002CT |
| mature neutrophil | Blueprint | C0010K | C000S5 |
| conventional dendritic cell | Blueprint | S00CP651 | S00D71 |
| adipose | ENCODE | ENCFF318AMC | ENCFF477GKI |
| HepG2 | ENCODE | ENCFF847OWL | ENCFF064GJQ |
| pancreas | ENCODE | ENCFF753ZMQ | ENCFF500DKA |
| small intestine | ENCODE | ENCFF266NGW | ENCFF122LEF |
| spleen | ENCODE | ENCFF550FZT | ENCFF333OHK |
| stomach | ENCODE | ENCFF435SPL | ENCFF497YOO |
| tibial nerve | ENCODE | ENCFF843SYR | ENCFF699KTW |
| skeletal muscle myoblast primary cell | ENCODE | ENCFF774GXJ | - |

#### 3 Supplementary figures

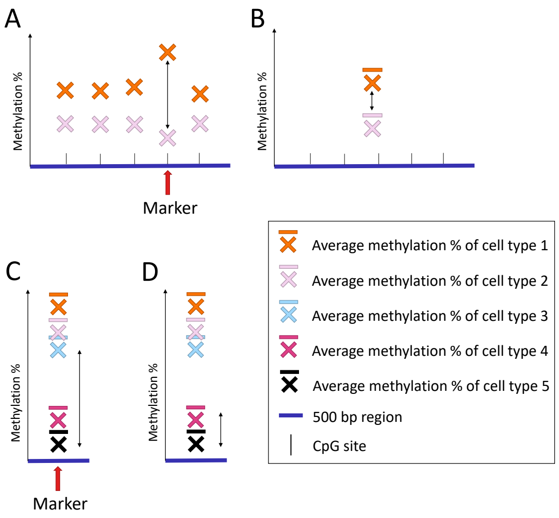


Fig. S1: Illustration of the two principal changes to the approach for findings markers in the genome. The arrows indicate the difference between cell types. Figures **(A)** and **(C)** illustrate how the markers are found originally, and Figures **(B)** and **(D)** how they are found after improvements. Figures **(A)** and **(B)** show how measuring the distance between single CpG sites **(A)** results in different markers than measuring the distance between 500 bp regions **(B)**. Cell types 1 and 2 do not have a large distance when regarding their average over the entire region, making this region an unsuitable marker. Figures **(C)** and **(D)** show that the distance from the median cell type **(C)** is different from the distance from the min cell type **(D)**. Using the median would result in a marker that does not differentiate well between cell types 4 and 5.

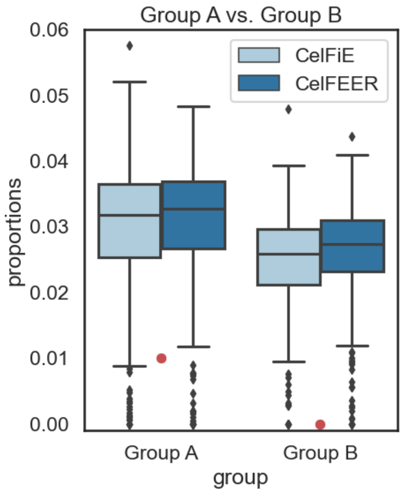


Fig. S2: Estimates of the proportion of a rare cell type (1%) that is present in group A but not in group B, estimated over 50 replicate runs using CelFiE and CelFEER. Only the estimated and true proportions of this rare cell type are plotted. The true proportions are represented by the red dots.

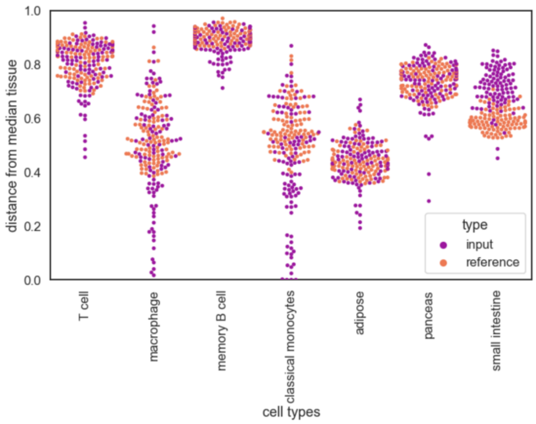


(a) CelFiE markers

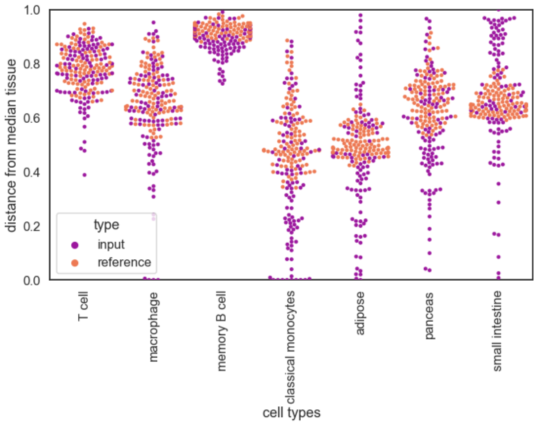


(b) CelFEER markers

Fig. S5: Markers found by **(a)** CelFiE and **(b)** CelFEER for seven different cell types.

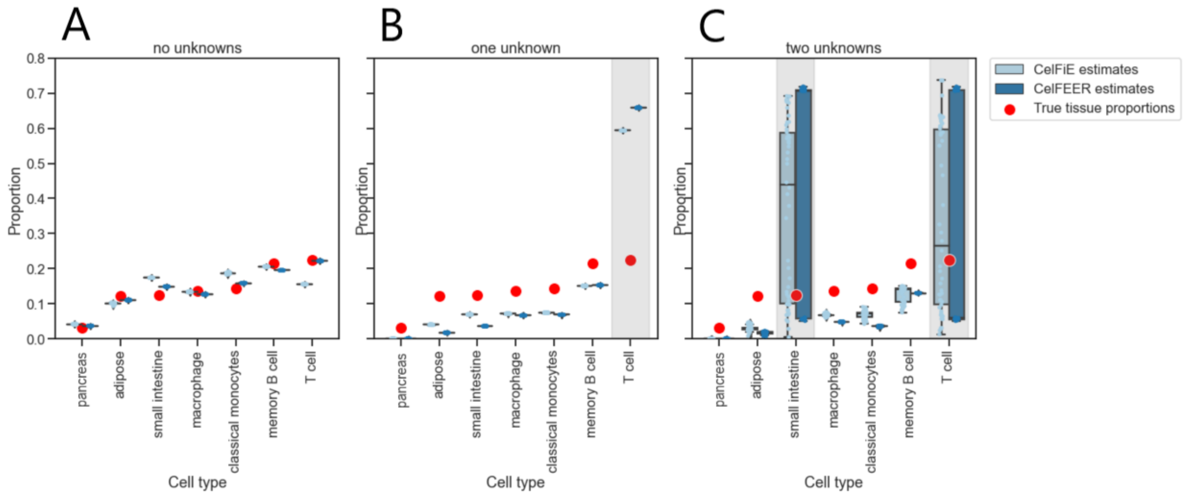


Fig. S3: Cell type proportions estimated by CelFiE and CelFEER for zero, one and two unknowns respectively. The boxplots visualize the estimated proportions of 50 replicates for a randomly chosen individual. On top of the boxplots, the individual datapoints are plotted.

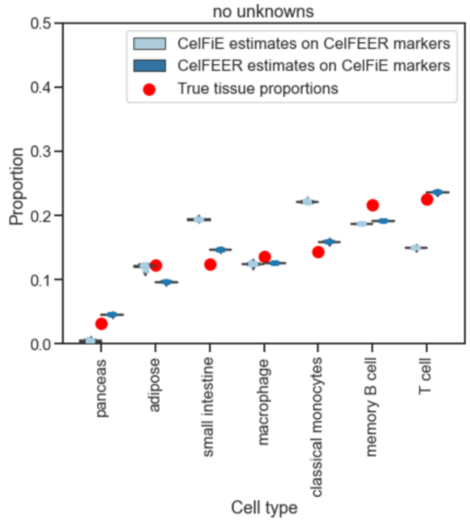


Fig. S4: CelFiE and CelFEER run on different markers.

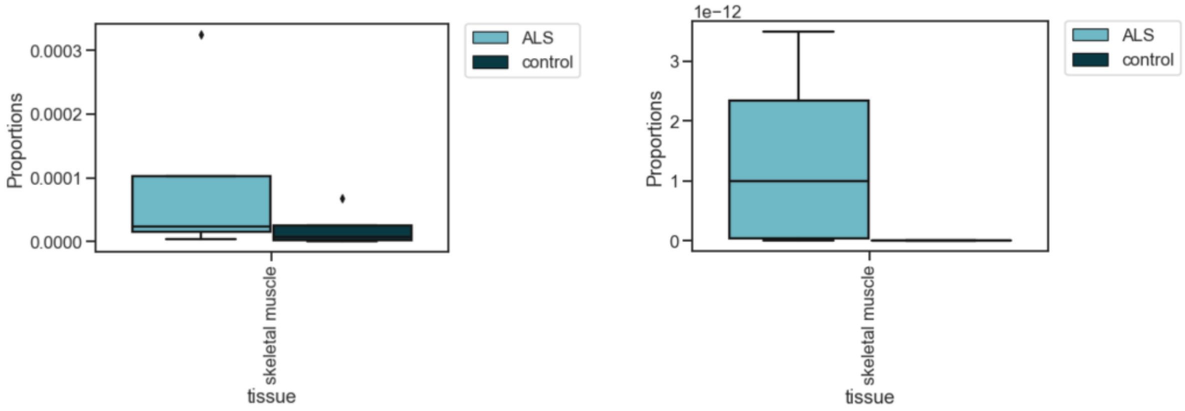


(a) CelFiE (b) CelFEER

Fig. S6: Estimated proportions of skeletal muscle cfDNA when one unknown cell type is estimated in addition to the total 19 cell types in the reference data.

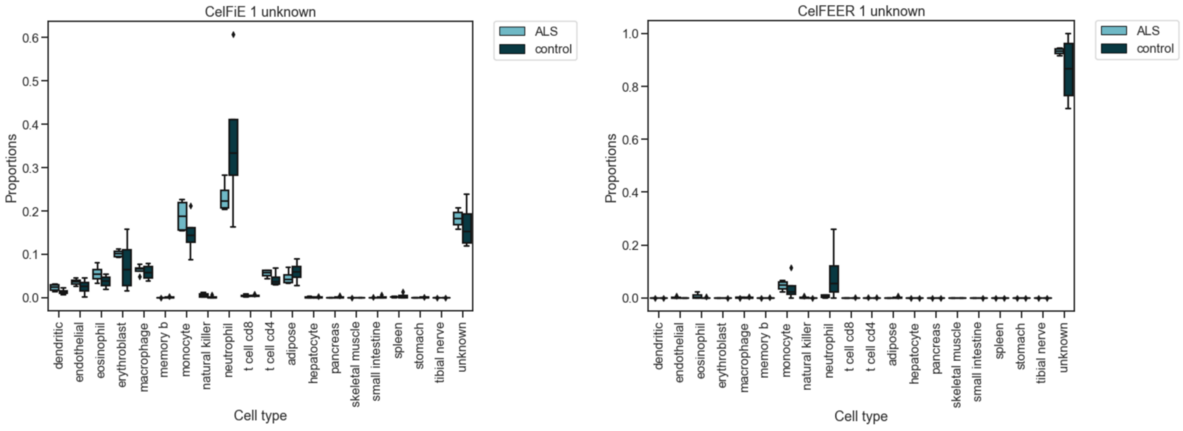


(a) CelFiE ALS and control cell type decomposition (b) CelFEER ALS and control cell type decomposition

Fig. S7: Complete cell type decomposition when one unknown cell type is estimated.

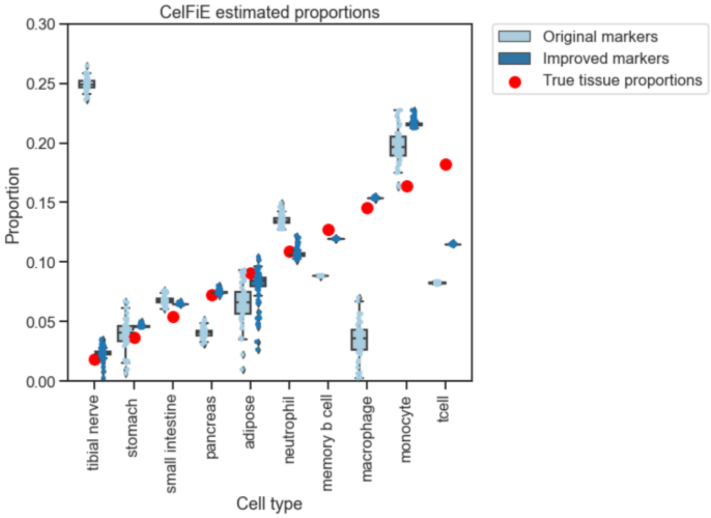


Fig. S8: CelFiE estimated cell type proportions on a simulated cfDNA mixture using WGBS cell type data, using both the markers found as described in [1] and the markers found using our improved method. For visualisation purposes the true cell type proportions are a simple incremental array summing to one. The results of 50 replicate runs on 10 individuals are displayed.

#### 4 Selection of cell type informative markers

A crucial step in predicting the cell type of origin is selecting markers in the genome that represent the cell types. Not only does a set of distinct markers improve prediction, it can make sequencing of cfDNA less expensive since only the DNA overlapping the markers needs to be sequenced. Methylation markers that span multiple CpG sites are in literature often referred to as differentially methylated regions. To find cell type informative markers, we started by analyzing the markers found using the method created by caggiano2021comprehensive, which is described in **??**. This method was then improved to find more informative markers. In this section and the following we refer to the absolute counts of methylated CpG sites as methylation values, and to the fraction of methylated to unmethylated CpG sites as methylation percentages.

##### Regions are more robust markers than single sites

caggiano2021comprehensive use the traditional approach of using single CpG sites as markers. This method, however, decreases the ability to differentiate between different cell types as it is sensitive to both biological and technical noise. In order to reduce noise, the CpG sites 250 bp upstream and 250 bp downstream of the markers are added to the markers' methylation counts. The authors showed that their method only returns sensible results when the methylation values are thus summed into regions. It nonetheless happens that the 500 bp surrounding the markers contain little CpG sites. This method does not exploit earlier findings that the methylation status is highly coupled between adjacent CpG sites [2]. Moreover, regions where CpG sites are clustered in high numbers, called CpG islands (CGIs), are known to be epigenetic regulatory regions that can be cell type specific [3].

According to these findings, it makes more sense to compare regions containing multiple CpG sites instead of single CpG sites to find differential markers. To test this hypothesis, CpG sites were grouped in a simple fashion: CpG sites were summed if they were in a 500 bp vicinity of each other. The starting location of each 500 bp window was set to be the first CpG site which contained measurements and did not fall in a previous bin. This strategy has the downside that it may split clusters in two, but if this is the case and if this cluster is differential, it is not harmful for the method to use both parts of the cluster as markers.

In addition to summing over 500 bp windows, we also summed over 10 bp windows with the idea of removing noise while still looking at mostly local methylation. After finding markers on the 10 bp windows, the surrounding CpG sites were summed to nevertheless obtain a total window of 500 bp. In order to compare the markers' ability to differentiate between cell types, we looked at the absolute difference between the methylation percentage of each marker's cell type and the median methylation percentage of all cell types. To test the generalizability of the markers, we did this for both the reference data (which was used to find the markers) and for the input data. As can be seen in Figure S9, the markers are most differential when they are first summed in 500 bp windows, and the variance in distance has substantially decreased. This strategy also seems to result in markers that generalize relatively well to unseen data, as the input and reference data have a similar distance to the median of other cell types. Although summing in 500 bp windows seems to return better markers than summing in 10 bp windows, it is remarkable how much improvement can be seen compared to the original method, especially for the tibial nerve cells. This is probably the effect of the decrease in noise which appears even if we sum over such small intervals. The results confirm the belief that markers are more differentiable when CpG sites are first summed compared to when they are summed after selecting individual sites. For this reason, all future experiments on markers are done on sites summed in 500 bp regions. In this section, we used only hypomethylated markers as they promised to be most distinguishing between cell types.

##### Hypomethylated sites are easier to differentiate than hypermethylated sites or than a mixture of both

caggiano2021comprehensive originally determined the best markers for each cell type by comparing the distances between the methylation percentages of each individual cell type to the median methylation percentage of all cell types. This should, in theory, result in a mixture of hypo- and hypermethylated markers. A sufficiently large distance to the median is, however, not a very strict requirement as it does not remove the probability of having two or more cell types with a very similar methylation percentage (especially as the number of cell types in the reference grows). Moreover, in practice almost all of the markers found using this method are hypomethylated, so there is little benefit in also allowing for hypermethylated markers.

To make the markers more differential, we measured the distance between the methylation percentage of each cell type and the minimum methylation percentage of all other cell types. This approach was compared to the original approach (where the distance from the median is measured instead) as well as to a similar approach where we looked only for hypermethylated markers (and thus compared to the maximum of all other cell types). When comparing the markers' distances from the median, the original method seems to result in the best markers for all cell types except adipose (Figure S9d). Hypomethylated markers, on the other hand, have a slightly smaller distance from the median for all cell types except for adipose, for which the distance is larger (Figure S9e). Hypermethylated markers have overall the smallest distance from the median (Figure S9c).

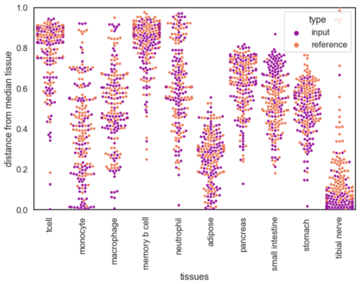


(a) Individual sites

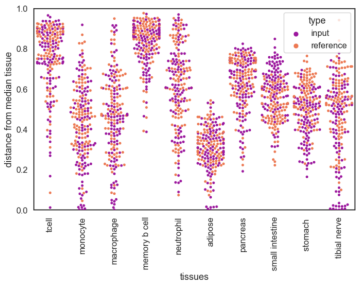


(b) Summed over 10 bp

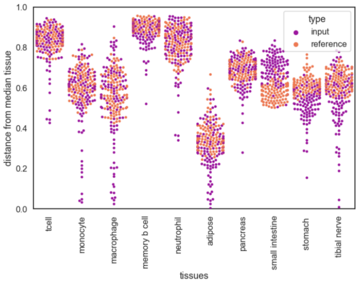


(c) Summed over 500 bp

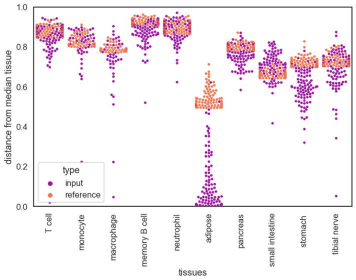


(d) Distance measured from median

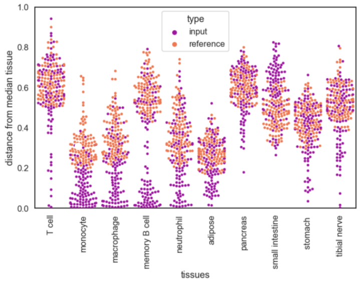


(e) Hypermethylated

Fig. S9: Distance from median methylation percentage for three different strategies; Purple dots represent the input at different marker locations and orange dots represent the reference at the same marker locations. The reference data was used to find the marker locations.
Row 1: Comparison between single CpG site markers which are summed with their 500 bp neighbouring sites (S9a), 10 bp markers which are summed with their 490 bp neighbouring sites (S9b) and 500 bp markers (S9c).
Row 2: Comparison between markers defined by their distance from the median methylation percentage (S9d), distance from the maximum (S9e) and distance from the minimum (S9c).
All figures in row one use hypomethylated markers, and all figures in row two are first summed over 500 bp.

However, as reasoned above, the distance from the median may not be the best metric for defining the ability to differentiate between cell types. Therefore, we can not assume that the distance from the median also translates to the best cell type deconvolution results. For this reason, we looked at the results on a simulated mixture of the WGBS data of 10 cell types and measured the Pearson's correlation between the true and estimated cell type proportions of 50 replicate runs for 10 individuals. We set the true cell type proportions to a linearly incrementing array that sums to one. While the hypomethylated markers resulted in a correlation of $r^2 = 0.86 \pm 0.01$, the hypermethylated and original method resulted in a correlation of $r^2 = 0.68 \pm 0.04$ and $r^2 = 0.58 \pm 0.03$ respectively. This confirms the idea that the distance from the median is not the best metric for obtaining differentiable markers.

This can additionally be observed from the amount of markers found by each metric. The method for finding markers works in such way that it first finds the 100 best markers for each cell type and then removes the markers that are overlapping multiple cell types. As can be seen in Figure S10, the original method finds less markers which means that the markers it finds have a high amount of overlap between cell types. Especially monocytes and macrophage cells seem to have much overlap, which makes sense given the fact that macrophage cells are differentiated monocyte cells [4]. Hypo- and hypermethylated markers are nevertheless able to differentiate these two cell types. To test whether the markers found using the original method would result in better performance if more markers were included, we first tested for uniqueness of the 200 best markers of each cell type and then included the 100 best markers. This way each cell type had 100 markers. This resulted in a negligible increase in performance.

As the hypomethylated markers seem to give the best results, all experiments in this section, including the previous section, use hypomethylated markers.

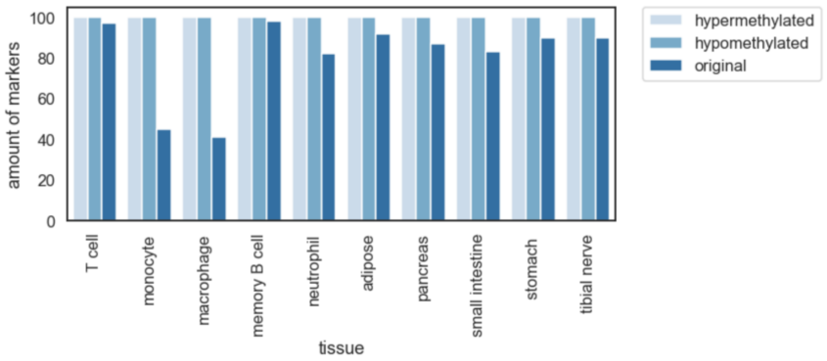


Fig. S10: The bar chart shows the amount of markers found for each cell type using each of the three different ways to measure the distance between cell types.

##### Additional improvements for increased differentiation between cell types

In addition to the improvements discussed in the previous two sections, there were two possible unwanted outcomes in the original method for finding markers. The first of which is that the authors introduced only a requirement for the median read depth of all cell types at a candidate marker site. This means that if one cell type is covered by one single read only at a candidate CpG site, this CpG site can still become a marker for that cell type as long as all other cell types have sufficient coverage. A simple adjustment was made to the method by setting a minimum depth threshold for cell types at their potential marker sites. This threshold was set equal to the median depth threshold.

The second possible undesirable behaviour is caused by the manner of checking for the uniqueness of the markers. As only the top 100 markers of all cell types is checked for overlapping markers, it is possible that the same site is the 100th best marker for cell type x and the 101st best marker for cell type y. This situation was prevented by keeping a list of the 150 best markers for each cell type which are all checked for uniqueness, such that the 100th best marker for cell type x could not even be the 150th best marker for cell type y.

The effects of both changes were measured by calculating the Pearson's correlation between the true and estimated cell type proportions for 10 individuals and 10 cell types of 50 replicate runs. The true cell type proportions were drawn from a uniform distribution and made to sum to one. Using no improvements, the correlation between the true and estimated cell types was $r^2 = 0.87 \pm 0.09$ . Using only the additional uniqueness criterion did not change the results, and resulted in the same amount of correlation. The stricter depth criterion, however, improved the correlation to $r^2 = 0.91 \pm 0.06$. Combining both improvements resulted in the same correlation. This means that the situation described above does not occur, and the markers are already sufficiently unique. This is perhaps a consequence of using hypomethylated markers only.

#### References

[1] Caggiano, C., Celona, B., Garton, F., Mefford, J., Black, B. L., Henderson, R., Lomen-Hoerth, C., Dahl, A., and Zaitlen, N. (2021) Comprehensive cell type decomposition of circulating cell-free DNA with CelFiE. *Nature communications,* **12**(1), 1–13.

[2] Guo, S., Diep, D., Plongthongkum, N., Fung, H.-L., Zhang, K., and Zhang, K. (2017) Identification of methylation haplotype blocks aids in deconvolution of heterogeneous tissue samples and tumor tissue-of-origin mapping from plasma DNA. *Nature genetics,* **49**(4), 635–642.

[3] Tahir, R. A., Zheng, D., Nazir, A., and Qing, H. (2019) A review of computational algorithms for CpG islands detection. *Journal of biosciences,* **44**(6), 1–11.

[4] Yang, J., Zhang, L., Yu, C., Yang, X.-F., and Wang, H. (2014) Monocyte and macrophage differentiation: circulation inflammatory monocyte as biomarker for inflammatory diseases. *Biomarker research,* **2**(1), 1–9.
